# An extended catalogue of tandem alternative splice sites in human tissue transcriptomes

**DOI:** 10.1101/2020.09.11.292722

**Authors:** Aleksei Mironov, Stepan Denisov, Alexander Gress, Olga V. Kalinina, Dmitri D. Pervouchine

**Affiliations:** Skolkovo Institute for Science and Technology, Moscow 143025, Russia; Helmholtz Institute for Pharmaceutical Research Saarland (HIPS), Helmholtz Centre for Infection Research (HZI), 66123 Saarbrücken, Germany; Faculty of Medicine, Saarland University, 66421 Homburg, Germany

## Abstract

Tandem alternative splice sites (TASS) is a special class of alternative splicing events that are characterized by a close tandem arrangement of splice sites. Most TASS lack functional characterization and are believed to arise from splicing noise. Based on the RNA-seq data from the Genotype Tissue Expression project, we present an extended catalogue of TASS in healthy human tissues and analyze their tissue-specific expression. The expression of TASS is usually dominated by one major splice site (maSS), while the expression of minor splice sites (miSS) is at least an order of magnitude lower. Among 73k miSS with sufficient read support, 12k (17%) are significantly expressed above the expected noise level, and among them 2k are expressed tissue-specifically. We found significant correlations between tissue-specific expression of RNA-binding proteins (RBP) and tissue-specific expression of miSS that is consistent with miSS response to RBP inactivation by shRNA. In combination with RBP profiling by eCLIP, this allowed prediction of novel cases of tissue-specific splicing regulation including a miSS in QKI mRNA that is likely regulated by PTBP1. According to the structural annotation of the human proteome, tissue-specific miSS are enriched within disordered regions, and indels induced by miSS are enriched with short linear motifs and post-translational modification sites. Nonetheless, more than 15% of tissue-specific miSS affect structured protein regions and may adjust protein-protein interactions or modify the stability of the protein core. The significantly expressed miSS evolve under the same selection pressure as maSS, while other miSS lack signatures of evolutionary selection and conservation. Using mixture models, we estimated that not more than 10% of maSS and not more than 50% of significantly expressed miSS are noisy, while the proportion of noisy splice sites among not significantly expressed miSS is above 70%.

## Introduction

Alternative splicing (AS) of most mammalian genes gives rise to multiple distinct transcript isoforms that are often regulated between tissues [1–3]. It is widely accepted that among many types of AS, exon skipping is the most frequent subtype [1]. The second most frequent AS type is the alternative choice of donor and acceptor splice sites, the major subtype of which are the so-called tandem alternative splice sites (TASS) that are located only a few nucleotides from each other [4, 5]. About 15–25% of mammalian genes possess TASS, and they occur ubiquitously throughout eukaryotes, in which alternative splicing is common [4]. TASS were experimentally shown to be functionally involved in DNA binding affinity [6], subcellular localization [7], receptor binding specificity [8] and other molecular processes (see [4] for review).

The outcome of the alternative splicing of non-frameshifting TASS on the amino acid sequence encoded by the transcript is equivalent to that of short genomic indels. The latter cause broad genetic variation in the human population and impact human traits and diseases [9, 10]. For example, cystic fibrosis is often caused by an indel that leads to a deletion of a single amino acid from the protein sequence [11]. For a different type of alternative splicing with a similar effect on amino acid sequence, alternative microexons, it has been demonstrated that insertion of two amino acids may influence protein-protein interactions in brains of autistic patients [12]. Structural analysis of non-frame-shifting genomic indels revealed that they predominantly adopt coil or disordered conformations [13]. Likewise, non-frame-shifting TASS with significant expression of multiple isoforms are overrepresented in the disordered protein regions and are evolutionarily unfavourable in structured protein regions [14].

The two most studied classes of TASS are the acceptor NAGNAGs [5, 15, 16] and the donor GYNNGYs [17]. In these TASS classes, alternative splicing is significantly influenced by the features of the cis-regulatory sequences, but very little is known about their function, tissue-specific expression, and regulation [5, 17, 18]. Recent genome-wide studies estimated that at least 43% of NAGNAGs and ∼20% of GYNNGYs are tissue-specific [5, 17]. It is believed that closely located TASS such as NAGNAGs and GYNNGYs originate from the inability of the spliceosome to distinguish between closely located cis-regulatory sequences, and therefore most TASS are attributed to splicing errors or noise [17, 19, 20]. However, it is not evident from the proteomic data what fraction of alternative splicing events and, in particular, of TASS splicing indeed lead to the changes in the protein aminoacid sequence [21–23].

The biggest existing catalogue of TASS, TASSDB2, is based on the evidence of transcript isoforms from expressed sequence tags [24]. The advances of high-throughput sequencing technology open new possibilities to identify novel TASS [25]. Here, we revisit the catalogue of TASS by analyzing a large compendium of RNA-seq samples from the Genotype Tissue Expression (GTEx) project [26]. We substantially extend the existing catalogue of TASS, characterize their common genomic features, and systematically describe a large set of TASS that have functional signatures such as evolutionary selection, tissue-specificity, impact on protein structure, and regulation. While it is believed that the expression of TASS primarily originates from splicing noise, here we show that a number of previously unknown TASS may have important physiological functions and estimate the proportion of noisy splicing of TASS.

## Results

### The catalogue of TASS

In order to identify TASS, we combined three sources of data. First, we extracted the annotated splice sites from GENCODE and NCBI RefSeq human transcriptome annotations [27, 28]. This resulted in a list of ∼570k splice sites, which will be referred to as annotated. Next, we identified donor and acceptor splice sites in split read alignments from the compendium of RNA-seq samples from the Genotype Tissue Expression Project (GTEx) [26] by pooling together its 8,548 samples. This resulted in a list of ∼1 mln splice sites, which will be referred to as expressed. A splice site may belong to both these categories, i.e. be annotated and expressed, or be annotated and not expressed, or be expressed and not annotated (the latter are referred to as *de novo*). Third, we took the list of intronic splice sites from [29] with MaxEntScan score greater than 3 and excluded ones that were previously called expressed or annotated. This resulted in a list of ∼3.47 mln sequences that are similar to splice sites, but have no evidence of expression or annotation and will therefore be referred to as cryptic. The combined list from all three sources contained ∼4.58 mln unique splice sites (S1 Table,A).

A TASS cluster is defined as a set of at least two splice sites of the same type (either donor or acceptor) such that each two successive splice sites are within 30 nts from each other (Fig 1,A). The number of splice sites in a TASS cluster will be referred to as the cluster size. According to this definition, each splice site can belong either to a TASS cluster of size 2 or larger, or be a standalone splice site. Among ∼4.58 mln candidate splice sites in the initial list, a large fraction (∼1.54 mln) belong to TASS clusters (S1 Table,B).

**Fig 1.**
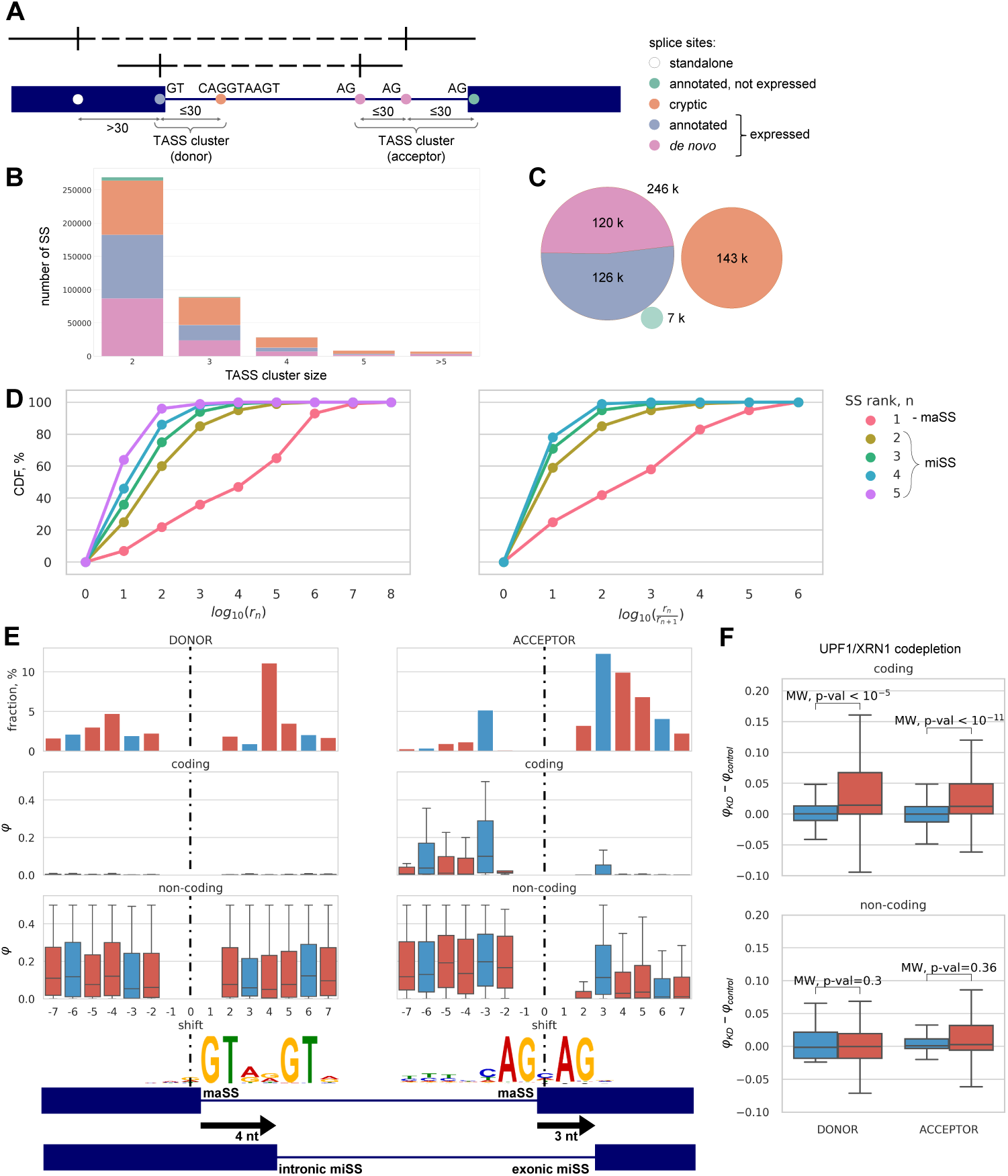
The catalogue of TASS. (A) Splice sites (SS) are categorized as annotated (GENCODE and Refseq), *de novo* (inferred from RNA-seq) or cryptic (detected by MaxEntScan). TASS clusters consist of splice sites of the same type (donor or acceptor) such that each two consecutive ones are within 30 nts from each other. (B) TASS cluster size distribution. (C) The number of annotated, de novo and cryptic TASS. (D) The expression of the major SS (maSS, i.e. rank 1) and minor splice sites (miSS, i.e. rank 2 or higher). Left: the cumulative distribution of *rn*, the number of split reads supporting maSS and miSS. Right: the cumulative distribution of *r*_*n*_ relative to *r*_*n*+1_. (E) The distribution of shifts, i.e., the positions of miSS relative to maSS (top) for miSS of rank 2. The relative usage of miSS (*φ*) in coding (middle) and non-coding regions (bottom). Logo chart of miSS sequences of +4 donor shifts and of +3 acceptor shifts. (F) The change of miSS relative usage (*φ*_*KD*_ *− φ*_*control*_) upon NMD inactivation for frame-preserving (blue) and frame-disrupting (red) miSS.

About 99% of splice sites in TASS clusters are located in clusters of size 2, 3, 4 and 5 (Fig 1,B). In what follows, we confined our analysis to TASS clusters consisting of 5 or fewer splice sites and having at least one expressed splice site (S1 Table,C). This way we obtained ∼396k splice sites; among them ∼126k expressed annotated splice sites, ∼120k expressed *de novo* splice sites, ∼7k annotated splice sites that are not expressed, and ∼143k cryptic splice sites (Fig 1,C). We categorized a TASS cluster as coding if it contained at least one non-terminal boundary of an annotated protein-coding exon, and non-coding otherwise. Most annotated TASS are coding while most *de novo* TASS are non-coding (S1 Fig).

Whereas almost a half of the expressed splice sites are *de novo* (Fig 1,C), their split read support is much lower than that of the annotated splice sites (Table 1). We pooled together the read counts from all 8,548 GTEx samples and ranked splice sites within each TASS cluster by the number of supporting reads (Fig 1,D, left). The dominating splice site (rank 1, also referred to as major splice site, or maSS) is expressed at a much higher level compared to splice sites of rank 2 or higher (referred to as minor splice sites, or miSS), and maSS in a TASS cluster is expressed several orders of magnitude stronger relative to all miSS (Fig 1,D, right). We identified the total of 72,652 expressed miSS, the majority of which (87%) were not annotated, unlike the expressed maSS, 67% of which were annotated (Table 2). At that, the set of expressed miSS significantly extends TASSDB2 database, which is limited to TASS separated by 2–12 nt and contains only 3,172 expressed miSS [24], as we found 32,415 such miSS that are absent from TASSDB2 (S2 Fig).

**Table 1.** Abundance and split read support of annotated and *de novo* TASS.

|  | expressed TASS |  | % of split reads supporting TASS |
| --- | --- | --- | --- |
|  | number | % |  |
| total | 245,976 | 100% | 100% |
| annotated | 125,891 | 51.18% | 99.89% |
| *de novo* | 120,085 | 48.82% | 0.21% |

**Table 2.** The fractions of annotated and *de novo* sites among maSS and miSS.

|  | *de novo* | annotated | total |
| --- | --- | --- | --- |
| maSS | 57,066 (33%) | 116,258(67%) | 173,324 |
| miSS | 63,019 (87%) | 9,633(13%) | 72,652 |

Previous large-scale analyses of human RNA-seq samples have noted that splice junctions called from the split reads mostly involve only one novel splice site, while only in a minor fraction of them neither donor nor acceptor splice site is annotated [30]. Indeed, we checked that only 2.2% of split reads that support miSS on one end support several splice sites on the other end. Therefore, to quantify the relative usage of a miSS, we introduced the metric *φ* that takes into account only one end of the split read. It is defined as the number of split reads supporting a miSS as a fraction of the combined number of split reads supporting miSS and maSS. In contrast to the conventional percent-spliced-in metric for exons [31], *φ* measures the expression of a miSS relative to that of the corresponding maSS and takes into account only one end of the supporting split read (S3 Fig).

Since each miSS is associated with a uniquely defined maSS within a TASS cluster, the position of the miSS relative to the position of the maSS, which will be referred to as shift, is defined uniquely. Positive shifts correspond to miSS located downstream of the maSS in a gene, while miSS with negative shifts are located upstream of the maSS. Consistent with previous observations [32], the distribution of shift values for miSS in TASS clusters of size 2 reveals that the most frequent shifts among donor miSS are ±4 nts, while acceptor miSS are most frequently shifted by ± 3 nts (Fig 1,E, top). These characteristic shifts likely arise from splice site consensus sequences, e.g. NAGNAG acceptor and GYNNGY donor splice sites [17, 33]. Donor miSS in TASS clusters of size larger than 2 are often separated by an even number of nucleotides, while acceptor miSS are often separated by a multiple of 3 nts. For example, rank 2 and rank 3 donor miSS are often located 2 or 4 nts from the maSS and tend to have shifts with opposite signs, while rank 2 and rank 3 acceptor miSS are often separated by 3 nts and tend to be located downstream of the maSS (S4 Fig). In what follows, we refer to miSS that are located outside of annotated exons as intronic, and exonic otherwise (Fig 1,E, bottom). Intronic miSS correspond to insertions, while exonic miSS correspond to deletions.

In the coding regions, we expect the distance between miSS and maSS to be a multiple of 3 in order to preserve the reading frame. Indeed, shifts by a multiple of 3 nts are the most frequent among coding acceptor miSS, however a considerable proportion of shifts by not a multiple of 3 nts also occur in coding donor and acceptor miSS (S5 Fig, top). We therefore asked whether these frame-disrupting shifts are actually expressed and found that, in spite of their high frequency, the relative expression of miSS in the coding regions, as measured by the *φ* metric, is still dominated by shifts that are multiple of 3 nts (Fig 1,E, middle), while the relative expression of miSS in the non-coding regions doesn’t depend on the shift (Fig 1,E, bottom). Consistent with this, frame-disrupting miSS in the coding regions are significantly upregulated after inactivation of the nonsense mediated decay (NMD) pathway by the co-depletion of two major NMD components, UPF1 and XRN1 [34], while no such upregulation is observed in the non-coding regions (Fig 1,F). This indicates that broad positional repertoire of frame-disrupting shifts in coding TASS is efficiently suppressed by NMD.

We next asked whether the expression patterns systematically differ for miSS located upstream and downstream of the maSS. In coding regions, the acceptor miSS are more often shifted downstream, while miSS located upstream tend to be expressed stronger than miSS located downstream (Fig 1,E). This observation supports splice junction wobbling mechanism, in which the upstream acceptor splice site is usually expressed stronger than the downstream one [18]. However, the expression difference can also be explained by subtle, yet systematic differences in splice site strengths. We found that miSS are on average weaker than maSS (S6 Fig,A), the strength of a miSS relative to the strength of maSS is correlated with its relative expression (S6 Fig,B), and the upstream acceptor miSS tend to be on average stronger than the downstream acceptor miSS (S6 Fig,C). Nonetheless, the upstream miSS are expressed stronger than the downstream miSS even for miSS that are similar in strength to their corresponding maSS (S6 Fig,D). For the acceptor splice sites, this finding supports the so-called linear scanning model, in which the spliceosome scans from the branch point towards the nearest acceptor splice site [35].

### Expression of miSS in human tissues

Tissue specificity is commonly considered as a proxy for splicing events to be under regulation [5, 36]. To assess the tissue-specific expression of miSS, we calculated the *φ*_*t*_ metric, i.e. the relative expression of miSS with respect to maSS, by aggregating GTEx samples within each tissue *t*. However, different tissues are represented by a different number of individuals and, consequently, TASS in different tissues have different read support. To account for this and for the dependence of the relative expression of miSS on the gene expression level [17, 37], we constructed a zero-inflated Poisson linear model that describes the dependence of miSS-specific read counts (*r*_*min*_) on maSS-specific read counts (*r*_*maj*_) (Fig 2,A). Using this model, we estimated the statistical significance of miSS expression and selected significantly expressed miSS using q-values with 5% threshold [38]. In what follows, we shortly refer to these significantly expressed miSS as significant.

**Fig 2.**
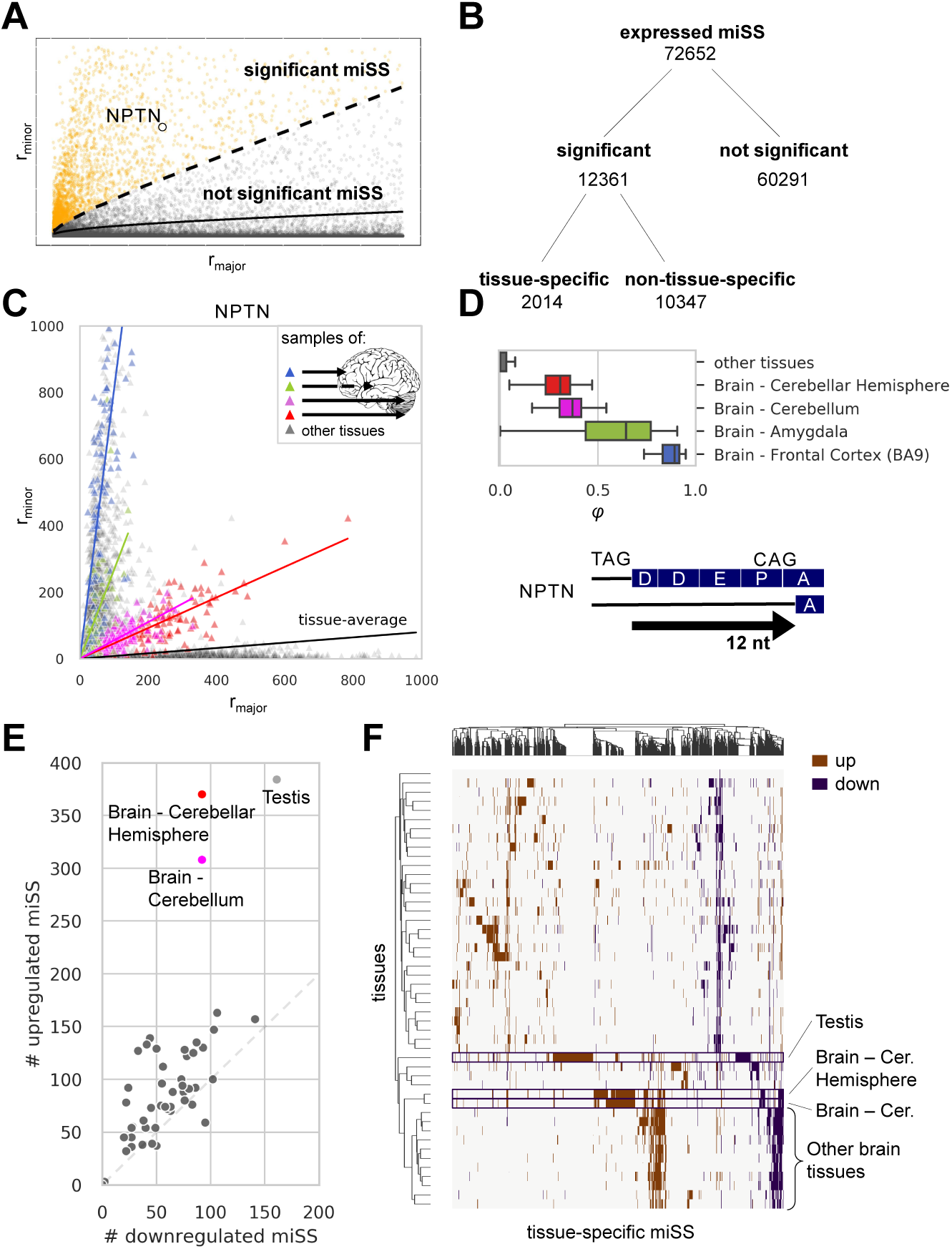
Expression of miSS in human tissues. (A) Zero-inflated Poisson model of miSS expression relative to maSS enables identification of significantly expressed miSS. Each dot represents a miSS. Significant miSS are colored orange (q-value*<* 5%), not significant miSS are colored grey (q-value *≥*5%). (B) The classification of expressed miSS. (C) Tissue-specific expression of a miSS in the NPTN gene. Each dot represents a sample, i.e. one tissue in one individual. Tissue specificity is estimated by a linear model with dummy variables corresponding to tissues. (D) The distribution of NPTN miSS *φ* values in selected tissues (top). The indel caused by NPTN miSS results in the deletion of DDEP motif from the aminoacid sequence (bottom). (E) The number of tissue-specific (up- or downregulated) miSS in each tissue. (F) The clustering of tissue-specific miSS and tissues based on *φ* values.

Out of 72,652 expressed miSS, 12,361 (17%) were significantly expressed in at least one tissue (Fig 2,B). To identify tissue-specific miSS among significant miSS, we built a linear model with dummy variables corresponding to each tissue (see Methods). A miSS was called tissue-specific if the slope of the dummy variable corresponding to at least one tissue was statistically discernible from zero (q-value<0.05), i.e. the proportion of reads supporting a tissue-specific miSS deviates significantly from the average across tissues. As a result, we obtained a conservative list of 2,014 tissue-specific miSS (Fig 2,B). In the coding regions, tissue-specific miSS preserve the reading frame more often than non-tissue-specific miSS do (S2 Table); they also have on average stronger consensus sequences and, among the latter, frame-preserving miSS have a stronger evidence of translation according to Ribo-Seq data (S7 Fig,A). The intronic region nearby tissue-specific and significantly expressed miSS tends to be more conserved evolutionarily compared to not significant miSS (S7 Fig,C), with frame-disrupting miSS being significantly less conserved than frame-preserving miSS (S7 Fig,D).

One notable example of a tissue-specific miSS is in the exon 7 of NPTN gene, which encodes neuroplastin, an obligatory subunit of Ca^2+^-ATPase, required for neurite outgrowth, the formation of synapses, and synaptic plasticity [39, 40]. The slope of the linear model has a distinct pattern of variation across tissues, and moreover within brain subregions (Fig 2,C). Brain-specific expression of the acceptor miSS instead of maSS in NPTN leads to the deletion of Asp-Asp-Glu-Pro (DDEP) sequence from the canonical protein isoform (Fig 2,D).

Tissues differ by the number of tissue-specific miSS and by the proportion of miSS that are upregulated or downregulated. The sign of the slope in the linear model that describes the dependence of *r*_*min*_ on *r*_*maj*_ allows to distinguish up- and downregulation. In agreement with previous reports on alternative splicing [41], a number of tissues including testis, cerebellum, and cerebellar hemisphere harbor the largest number of tissue-specific miSS (Fig 2,E, S3 Table). The testis and the brain have a distinguished large set of miSS with almost exclusive expression in these tissues that set them apart statistically from the other tissues (Fig 2,F).

A special class of TASS are the so-called NAGNAG acceptor splice sites, i.e., alternative acceptor sites that are located 3 bp apart from each other [33]. According to the current reports, they are found in 30% of human genes and appear to be functional in at least 5% of cases [15]. Here, we identified an extended set of 7,414 expressed acceptor miSS, of which 429 are tissue specific, that are located ±3 nt from maSS (S8 Fig,A) which reconfirms 92% of 1,884 alternatively spliced and 33.5% of 1,338 tissue-specific NAGNAGs reported by Bradley et al [5]. Furthermore, we identified 111 tissue-specific NAGNAGs that are not present in the previous lists [5]. Among them there is a NAGNAG acceptor splice site in the exon 20 of the MYRF gene, which encodes a transcription factor that is required for central nervous system myelination. The upstream NAG is upregulated in stomach, uterus and adipose tissues and downregulated in brain tissues (S8 Fig,B). Similarly, we identified an extended set of 3,873 expressed GYNNGY donor splice sites, i.e., alternative donor splice sites that are located 4 bp apart from each other (S8 Fig,C). This set reconfirms 54% of 796 GYNNGY donor splice sites reported by Wang et al [17]. Additionally, we identified 46 novel tissue-specific GYNNGYs including a donor splice site in the exon 2 of the PAXX gene (S8 Fig,D), the product of which plays an essential role in the nonhomologous end joining pathway of DNA double-strand break repair [42]. Unlike NAGNAGs, alternative splicing at GYNNGYs disrupts the reading frame and is expected to generate NMD-reactive isoforms [17].

The expression of splicing factors partially explains tissue-specific patterns of alternative splicing [43]. In order to identify the potential regulatory targets of splicing factors among miSS, we analyzed the data on shRNA depletion of 181 RNA-binding proteins (RBP) followed by RNA-seq and compared it with tissue-specific expression of miSS and RBP [44]. Our strategy was to identify tissues with significant up- or downregulation of a miSS responding to the inactivation of a splicing factor with the same signature of tissue-specific expression.

To this end, we identified miSS that are up- or downregulated upon RBP inactivation by shRNA-KD and matched them with the list of tissue-specific miSS and the list of differentially expressed RBP. As a result, we obtained a list of miSS-RBP-tissue triples (2,014 miSS x 103 RBP x 51 tissues, see Methods) that were characterized by three parameters, Δ*φ*_*t*_, Δ*RBP*_*t*_, and Δ*φ*_*KD*_, where Δ*φ*_*t*_ is the change of the miSS relative usage in the tissue *t*, Δ*RBP*_*t*_ is the change of the RBP expression in the tissue *t*, and Δ*φ*_*KD*_ is the response of miSS to RBP inactivation by shRNA-KD (Fig 3,A). We classified a miSS-RBP-tissue triple as co-directed if the correlation between RBP and miSS expression was concordant with the expected direction of miSS expression changes from shRNA-KD (e.g., if Δ*φ*_*t*_ *>* 0, Δ*RBP*_*t*_ *>* 0 and Δ*φ*_*KD*_ *<* 0) and anti-directed otherwise (Fig 3,B). That is, in co-directed triples the direction of regulation from the observed correlation and from shRNA-KD coincide, and in anti-directed triples they are opposite.

**Fig 3.**
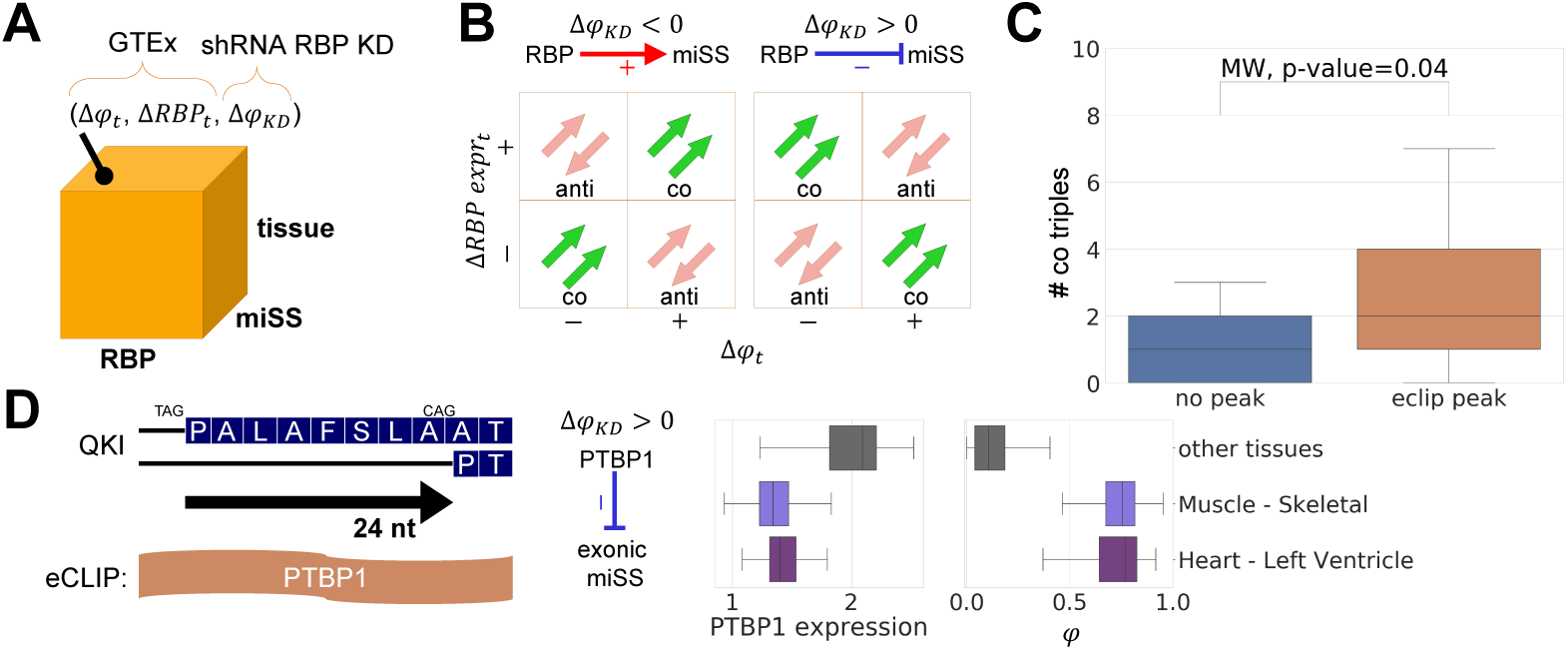
Regulation of miSS by RBP. (A) Each triple (miSS, RBP, tissue) is characterized by three metrics: (Δ*φt*, the change of miSS relative usage in tissue *t*; Δ*RBP*_*t*_, the change of the RBP expression in tissue *t*; and Δ*φ*_*KD*_, the change of miSS relative usage upon inactivation of RBP by shRNA-KD. (B) The response of miSS to RBP inactivation defines activating (Δ*φ*_*KD*_ *<* 0, red) and repressing (Δ*φ*_*KD*_ *>* 0, blue) regulation, which together with other metrics define co-directed and anti-directed triples. (C) The number of codirected miSS-RBP-tissue triples is significantly greater among triples supported by an eCLIP peak of the RBP near miSS as compared to non-supported triples. (D) A deletion of eight aminoacids in the QKI gene caused by the exonic miSS overlapping with PTBP1 eCLIP peak (left). The expression of PTBP1 (log_2_ *T P M*) is suppressed, while the miSS relative usage (*φ*) is promoted in muscle and heart (right). The miSS is activated in response to PTBP1 inactivation, suggesting its downregulation by PTBP1.

In order to obtain a stringent list of predictions, we applied 5% FDR threshold correcting for testing in multiple tissues, RBP, and miSS, and additionally required that miSS relative usage and RBP expression change not only significantly, but also substantially (|Δ*φ*_*t*_| *>* 0.05, |Δ*φ*_*KD*_| *>* 0.05, and |Δ*RBP*_*t*_| *>* 0.5. As a result, we obtained 138 co-directed and 93 anti-directed miSS-RBP-tissue triples. The number of co-directed triples is significantly greater than the number of anti-directed triples (binomial test, p-value=0.004). Next, we compared our predictions to the footprinting of RBP by the enhanced crosslinking and immunoprecipitation (eCLIP) method [45]. We found that miSS-RBP pairs with an eCLIP peak in the vicinity of the miSS tend to be more co-directed in comparison with miSS-RBP pairs without a eCLIP peak (Fig 3,C). Using eCLIP, we identified the total of 7 miSS-RBP candidate pairs of tissue-specific splicing regulation supported by an eCLIP peak (S5 Table). For example, the upregulation of an exonic acceptor miSS in exon 6 of the QKI gene in muscle and cardiac tissues is likely mediated by PTBP1 (Fig 3,D), consistent with previous findings that QKI and PTBP1 coregulate alternative splicing during muscle cell differentiation [46].

### Structural annotation of miSS

Alternative splicing of non-frameshifting TASS results in mRNA isoforms that translate into proteins with only a few amino acids difference. It was reported earlier that alternative splicing tends to affect intrinsically disordered protein regions [47], and that TASS with significant support from ESTs and mRNAs (506 such splice sites in total) are further overrepresented within regions lacking a defined structure [14].

We analyzed the structural annotation of the human proteome (see Methods) and found that significantly expressed miSS preferentially affect disordered protein regions, and tissue-specific miSS are found in disordered regions even more frequently (Fig 4,A, S10 Fig,B). We found that indels within disordered protein regions that are caused by TASS are enriched with short linear motifs (SLiMs), short sequence segments that often mediate protein interactions playing important functional roles in physiological processes and disease states [48–50] (Fig 4,B). Furthermore, we observed a significantly higher proportion of SLiMs related to post-translational modifications (PTM) in the indels caused by tissue-specific miSS (S10 Fig,C). The four most frequent PTM classes (acetylation, methylation, ubiquitination, phosphorylation) from the dbPTM database [51] are significantly enriched in protein sequences around miSS and in their nearby exonic regions for tissue-specific miSS (Fig 4,C). Interestingly, we found that nucleotide sequences of indels caused by tissue-specific miSS in disordered protein regions are more conserved evolutionarily compared to those of non-tissue-specific miSS, further supporting the enrichment of functional regulatory sites such as SLiMs or PTM (Fig 4,D).

**Fig 4.**
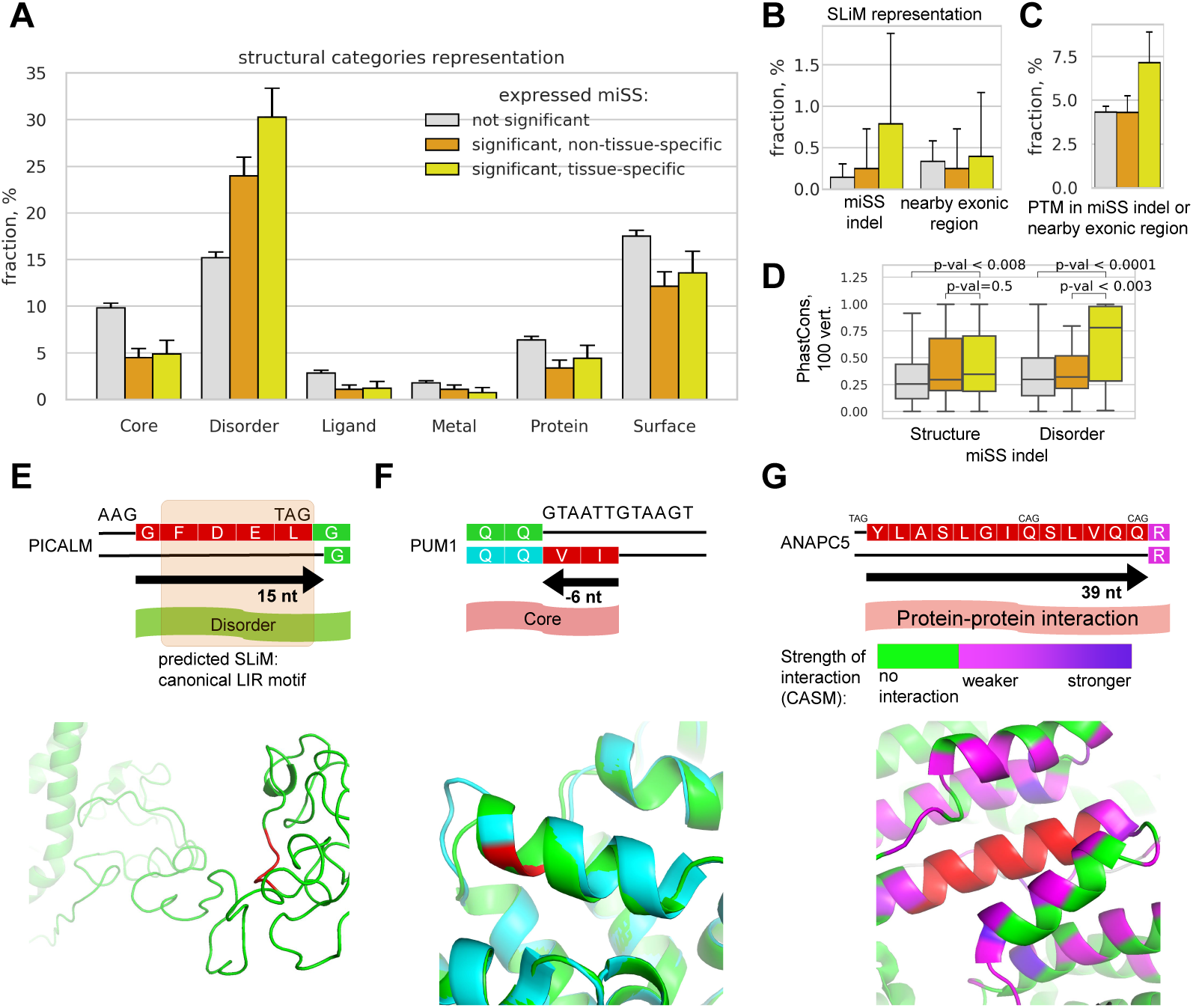
Structural annotation of miSS. **(A)** The proportion of miSS in genomic regions corresponding to protein structural categories. **(B)** The proportion of miSS in exonic regions corresponding to short linear motifs (SLiMs) from the ELM database and in nearby exonic regions (left). **(C)** The proportion of frame-preserving miSS in genomic regions corresponding to post-translational modification (PTM) sites from the dbPTM database. **(D)** The distribution of the average PhastCons conservation score (100 vertebrates) in the genomic regions between miSS and maSS. **(E)** The expression of an acceptor miSS in the predicted disordered region in the PICALM gene results in the deletion of five amino acids containing a predicted canonical LIR motif. The maSS-expressing structure of PICALM was modelled with I-TASSER (green); miSS indel is shown in red. **(F)** The expression of a donor miSS in the PUM1 gene results in the deletion of two amino acids from the core. The miSS-expressing structure is accessible at PDB (green, PDB ID: 1m8x); the maSS-expressing structure was modelled (cyan) and aligned to the miSS-expressing structure with I-TASSER. **(G)** The expression of an acceptor miSS in the ANAPC5 results in the deletion of 13 amino acids involved in protein-protein interactions. The maSS-expressing structure along with the interacting proteins is accessible at PDB (green, PDB ID: 6TM5); the miSS indel is shown in red. Computational alanine scanning mutagenesis (CASM) in BAlaS [57] was used to identify residues of the neighbouring proteins that contribute to the free energy of the interaction with the miSS indel region. The strength of the interaction (the positive change of the energy of interaction) is shown by the gradient color.

A notable example of a SLiM within indel that is caused by miSS is located in the PICALM gene, the product of which modulates autophagy through binding to ubiquitin-like LC3 protein [52, 53] (Fig 4,E). The expression of the short isoform lacking 15 nts at the acceptor splice site results in the deletion of Phe-Asp-Glu-Leu (FDEL) sequence, which represents a canonical LIR (LC3-interacting region) motif [54]. This motif interacts with LC3 protein family members to mediate processes involved in selective autophagy. This miSS is slightly upregulated in whole blood and downregulated in brain tissues consistently with a possible role in physiological regulation of autophagy.

Despite the enrichment of tissue-specific miSS in disordered regions, a sizable fraction of them (more than 10%) still correspond to functional structural categories such as protein core, sites of protein-protein interactions, ligand-binding, or metal-binding pockets (Fig 4,A). We therefore looked further into particular cases to discover novel functional miSS. For example, a shift of the donor splice site by 6 nts in the exon 17 of PUM1 gene results in a deletion of two amino acids (Fig 4,F). Only the structure of the miSS-expressing isoform of PUM1 is accessible in PDB (PDB ID: 1m8x) [55], and in it the site of deletion maps into an alpha helix. We modelled the structure of maSS-expressing isoform using I-TASSER web server [56]. The alignment of structures corresponding to the miSS- and maSS-expressing isoforms shows that the overall structure of the alpha helix can however be preserved, and only its raster is shifted by two amino acids. The insertion in the maSS-expressing isoform is shifted in this case into the preceding loop. The residues in this part of the helix become more hydrophobic, which may influence the overall helix or protein stability. Interestingly, this miSS is upregulated in skin, thyroid, adrenal glands, vagina, uterus, ovary, and testis, but downregulated in almost all brain tissues.

The expression of the miSS in exon 10 of ANAPC5 gene results in a 13 amino acids deletion from a protein interaction region (Fig 4,G). We modelled the interaction of these 13 amino acids with the adjacent protein structures using the computational alanine-scanning mutagenesis (CASM) in BAlaS [57]. We found 58 residues (49 residues in the ANAPC5 protein and 9 residues in the ANAPC15 protein), which, when mutated to alanine, cause a positive change in the energy of interaction with the 13 amino acid miSS indel region. The miSS is expressed concurrently with the maSS except for the brain tissues, in which the miSS is significantly downregulated. This may indicate the role of the miSS in various pathways in which ANAPC5 is involved as an important component of the cyclosome [58, 59].

In order to visualize structural classes associated with TASS, we created a track hub supplement for the Genome Browser [60]. The hub consists of three tracks: location of TASS indels, structural annotation of a nearby region, and tissue specific expression of selected TASS (S13 Fig). The catalogue of expressed miSS is also available in the table format (S6 Table).

### Evolutionary selection and conservation of miSS

In order to measure the strength of evolutionary selection acting on significantly expressed and tissue-specific miSS, and to evaluate how it compares with the evolutionary selection acting on maSS and splice sites outside TASS clusters, we applied a previously developed test for selection on splice sites [61]. We reconstructed the genome of a human ancestor taking marmoset and galago genomes as a sister group and an outgroup, respectively (see Methods). Using canonical consensus sequences of constitutive splice sites, we classified each nucleotide variant at each position as either consensus (Cn) or non-consensus (Nc) nucleotide (S11 Fig,AB). Then, we compared the frequency of Cn-to-Nc (or Nc-to-Cn) substitutions at different positions relative to the splice site (observed) with the background frequencies of the corresponding substitutions in neutrally-evolving intronic regions (expected) (S11 Fig,C). The ratio of observed to expected (*O/E*) equal to one indicates neutral evolution (no selection); *O/E >* 1 indicates positive selection; *O/E <* 1 indicates negative selection.

In the coding regions, the strength of negative selection acting to preserve Cn nucleotides in significantly expressed miSS is comparable to that in maSS and in constitutive splice sites, while no statistically discernible negative selection was detected in miSS that are not significantly expressed (Fig 5,A, left). A similar, yet weaker pattern of negative selection was observed for miSS in non-coding regions (S11 Fig,E, left). In contrast, the strength of positive selection, i.e., the *O/E* ratio for substitutions that create Cn nucleotides, is not significantly different from 1 in all miSS regardless of their expression, while a significant positive selection was detected in maSS and constitutive splice sites (Fig 5,A, right). This indicates that the evolutionary selection may preserve the suboptimal state of miSS relative to its corresponding maSS.

**Fig 5.**
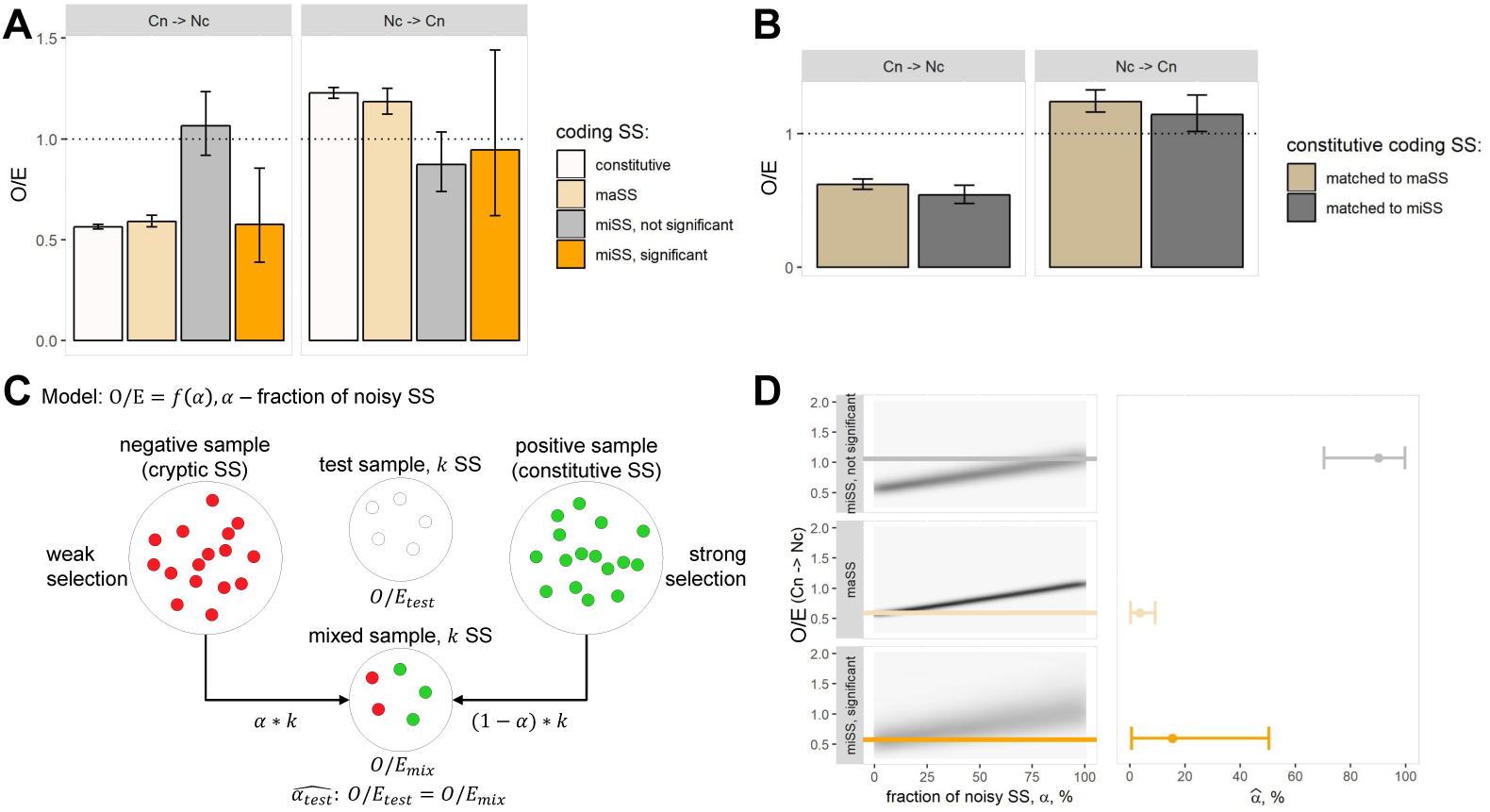
Evolutionary selection of miSS. **(A)** The strength of the selection, defined as the ratio of the observed (*O*) to the expected (*E*) number of substitutions, in selected categories of splice sites in the coding regions. The neutral expectation (*O/E* = 1) is marked by a dashed line. **(B)** The strength of the selection acting on constitutive coding splice sites matched to miSS and maSS by the ancestral splice site strengths. **(C)** The mixture model for the estimation of the fraction of noisy splice sites (*α*) using *O/E* ratio. A test sample of size *k* is modelled as a mixture of *αk* purely noisy (cryptic) splice sites and (1 *− α*)*k* purely functional constitutive splice sites. **(D)** The estimation of the fraction of noisy splice sites (*α*) from the observed values of *O/E*. Bootstrapped joint distribution of *O/E* and *α* values (left). 2.5%, 50% and 97.5% quantiles of the estimated *α* (right).

It was shown previously that the strength of the consensus sequence impacts evolutionary selection acting on a splice site [62, 63]. Indeed, the comparison of ancestral consensus sequences showed that constitutive splice sites and maSS have similar ancestral strengths, while miSS are considerably weaker (S11 Fig,F). To control for the influence of the splice site strength on the evolutionary selection, we sampled constitutive splice sites matching them by the ancestral strength with maSS and with miSS (Fig 5,B). However, despite a considerable difference in strengths, we observed only a subtle difference in evolutionary selection between constitutive splice sites that were matched to maSS and to miSS, indicating that the observed difference in selection acting on miSS and maSS is not due to weaker consensus sequences of miSS.

The difference in evolutionary selection between significantly expressed and the rest of miSS could arise from the difference in the fraction of noisy splice sites in these miSS categories. To estimate the fraction of noisy splice sites (*α*), we constructed a mixture model (Fig 5,C), in which we combined *αk* splice sites from the negative set of cryptic splice sites and (1 − *α*)*k* splice sites from the positive constitutive set and measured the strength of evolutionary selection in the combined sample for all values of *α*. Using this model, we constructed the joint distributions of *α* and *O/E* values of Cn-to-Nc substitutions for maSS, significantly expressed miSS, and the rest of miSS (Fig 5,D, left). From these distributions, we estimated 95% confidence intervals for the values of *α* that correspond to the actual *O/E* values in the observed samples (Fig 5,D, right; S11 Fig,G). The resulting estimates for the fraction of noisy splice sites among maSS, significantly expressed miSS, and the rest of the miSS are *<*10%, *<*50%, and *>*70%, respectively, indicating that at least a half of significantly expressed miSS are statistically discernible from noise.

## Discussion

Increasing amounts of high-throughput RNA-seq data have uncovered the expanding landscape of human alternative splicing [79]. Here, we present the most complete up-to-date catalogue of 72,652 miSS, of which 12,361 are significantly expressed in healthy human tissues according to GTEx data. It significantly extends previous catalogues constructed based on the evidence from expressed sequence tags [24] and also adds data on specific classes of TASS such as NAGNAGs [5] and GYNNGYs [17]. At the scale of RNA-seq data from GTEx, the sensitivity of detecting TASS must be high since the fraction of detected TASS is reaching a plateau with increasing the number of samples up to 8548 (S12 Fig). On the other hand, a substantial fraction of TASS are noisy (less than 50% among significantly expressed miSS and more than 70% among miSS that are not significantly expressed) reflecting natural tradeoff between sensitivity and specificity.

While the majority of miSS in the coding regions are located downstream of their respective maSS, the upstream miSS tend to be expressed stronger (Fig 1,E). That is, in spite of a broad repertoire of suboptimal splice sites near the major splice site, which are numerous but almost never used, the spliceosome tends to systematically choose a miSS that is located upstream, which leads to higher abundance of transcripts. This pattern could be a consequence of the linear scanning mechanism, in which the spliceosome traverses the pre-mRNA in the 5’ to 3’ direction. The relative expression is the strongest for the frame-preserving acceptor miSS, in support of the observation that transcript isoforms with frame-disrupting miSS are suppressed by NMD (Fig 1,F). We therefore expected that frame-disrupting miSS would be rare among significantly expressed and tissue-specific miSS, however almost a half of tissue-specific coding miSS disrupt the reading frame (S2 Table). Furthermore, frame-disrupting tissue-specific miSS are more conserved than non-significantly expressed miSS (S7 Fig,D) indicating a potential function such as, for example, fine-tuning of gene expression via NMD similar to the autoregulation of some RBPs by poison and essential exons [80–82].

Previous reports indicated that strongly expressed miSS located at a distance of 3, 6, 9 nt from the maSS in coding regions, which are to a large extent equivalent to the significantly expressed miSS introduced here, are overrepresented in disordered protein regions, with a more pronounced effect among acceptors than among donors [14]. Furthermore, the evolutionary selection against alternatively spliced NAGNAGs is strong in protein-coding regions and, in the latter, it is stronger in structured regions than in disordered regions [14]. Here, we reconfirmed this result and showed further that tissue-specific miSS are even more enriched in disordered protein regions, with the most pronounced effect being among acceptor exonic miSS, i.e. ones leading to the deletion in the protein sequence, and that tissue-specific miSS are associated with the presence of SLiMs and post translational modification sites. While there is no positive selection for Cn nucleotides among neither significantly expressed nor other miSS, we observed a strong negative selection acting to preserve Cn nucleotides in significantly expressed miSS (Fig 5,A). This finding can be explained by the tendency of functional miSS to preserve the suboptimal state relative to maSS, i.e functional miSS are evolutionarily conserved and maintain their Cn nucleotides, but they also do not harbor more Cn nucleotides not to outcompete maSS. Furthermore, we showed a tendency of many tissue-specific miSS to be regulated by RBP, e.g., the miSS in the exon 6 of the QKI gene is likely regulated by PTBP1 (Fig 3,D). All these findings are indicative of a functional role of at least a proportion of miSS.

The observations made in this manuscript are based on the analysis of RNA-seq data from the GTEx project [26]. It is hence worthwhile to address the question which proportion of these alternative splicing events translate to the protein level. Direct measurement of this proportion by, for example, shotgun proteomics is not instructive for many reasons, including limited coverage and low sensitivity of such experiments [83], as well as the fact that the cleavage site consensus of a widely used trypsin protease overlaps with the amino acid sequence induced by the splice site consensus, thus producing non-informative peptides [84]. This question has been debated in the literature [21–23]. On the one hand, proteomics data support expression of predominantly alternative splicing events such as mutually exclusive homologous exons [21], and alternative exons are under lower selection pressure [22]. On the other hand, ribosome profiling suggests expression of alternative isoforms [85], and functional studies on protein-protein interactions and signalling network rewiring indicate their functional importance [86]. Our study adds to this debate in that we have collected multiple lines of evidence that support expression on the protein level and functional importance of TASS-related isoforms. Our estimate of significantly expressed miSS largely exceeds the conservative estimate of proteomics-supported alternative splicing events [21]. Our analysis of Ribo-Seq experiments supports their expression, and in many cases this expression is tissue-specific. We also showed that significantly expressed miSS, as well as maSS, are under negative selection pressure. Finally, our analysis confirms that sites in protein sequence that correspond to TASS events are depleted from structured protein regions, just as for alternative splicing events in general [47, 87], which also suggests their non-neutral evolution and hence existence on the protein level. In line with previous research [87], we demonstrated that when located in disordered protein regions, TASS-associated events often affect sites of post-translational modification.

## Conclusion

Tandem alternative splice sites (TASS) are the second most abundant subtype of alternative splicing. The analysis of a large compendium of human transcriptomes presented here has uncovered a large and heterogeneous dataset of TASS, of which a significant fraction are expressed above the noise level and have signatures of tissue-specificity, evolutionary selection, conservation, and regulation by RBP. This suggests that the number of functional TASS in the human genome may be larger than it is currently estimated from proteomic studies despite the majority of TASS may represent splicing noise. The comprehensive catalogue of human TASS is available through a track hub for the Genome Browser (https://raw.githubusercontent.com/magmir71/trackhubs/master/TASShub.txt).

## Materials and Methods

### The catalogue of TASS

### The annotated splice sites

Throughout this paper, we use GRCh37 (hg19) assembly of the human genome which was downloaded from the UCSC genome browser [64]. To identify the annotated splice sites, we extracted internal boundaries of non-terminal exons from the comprehensive annotation of the GENCODE database v19 [27] and from UCSC RefSeq database [28]. As a result, we obtained 569,694 annotated splice sites (S1 Table,A).

### Expressed splice sites

The RNA-seq data from 8,548 samples in the Genotype-Tissue Expression (GTEx) consortium v7 data was analyzed as before [26]. Short reads were mapped to the human genome using STAR aligner v2.4.2a by the data providers [65]. Split reads supporting splice junctions were extracted using the IPSA package with the default settings [31] (Shannon entropy threshold 1.5 bit). At least three split reads were required to call the presence of a splice site. Samples of EBV-transformed lymphocytes and transformed fibroblasts and three samples with aberrantly high number of split reads were excluded. Only split reads with the canonical GT/AG dinucleotides were considered. Germline polymorphisms (SNPs, deletions and insertions) located within the splice site or within 35 nt of adjacent exonic regions were identified. Splice sites that were expressed exclusively in the samples, in which a polymorphism was present but absent in the other samples, were excluded to avoid split read misalignment caused by the discrepancy between the reference genome and the individual genotypes. This filtration removed 1.15% of expressed splice sites that were supported by 0.3% of the total number of split reads. As a result, we obtained 1,048,373 expressed splice sites (S1 Table,A).

### Cryptic splice sites

The intronic decoy splice sites from [29] were used as cryptic splice sites. Splice sites that were previously called expressed or annotated were excluded resulting in a list of 3,469,622 cryptic splice sites (S1 Table,A).

### Categorization of splice sites within TASS clusters

A TASS cluster (and all splice sites within it) was categorized as coding if it contained at least one non-terminal boundary of a coding exon, and non-coding otherwise. Thus, non-coding splice sites are located in untranslated regions (UTRs) of protein-coding genes or in other gene types such as long non-coding RNA. Splice sites were ranked based on the total number of supporting split reads. The splice site strength was assessed by MaxEntScan software [66] which computes a similarity of the splice site sequence and the consensus sequence. The higher MaxEnt scores correspond to splice site sequences that are closer to the consensus.

### Response of TASS clusters to NMD inactivation

To assess the response of TASS clusters to the inactivation of NMD, we used RNA-seq data from the experiments on co-depletion of UPF1 and XRN1, two key components of the NMD pathway [34]. Short reads were mapped to the human genome using STAR aligner v2.4.2a with the default settings. The read support of splice sites was called by IPSA pipeline as before (see processing of GTEx data). TASS in which the major splice site was supported by less than 10 reads were discarded. The response of a miSS to NMD inactivation was measured by *φ*_*KD*_ − *φ*_*C*_, where *φ*_*KD*_ is the relative expression in KD conditions and *φ*_*C*_ is the relative expression in the control.

### Expression of miSS in human tissues

#### Significantly expressed (significant) miSS

The number of reads supporting a splice site can be used for presence/absence calls, however it depends on the local read coverage in the surrounding genomic region and on the total number of reads in the sample [37, 67]. A good proxy for these confounding factors is the number of reads supporting the corresponding maSS. We therefore quantified the expression of miSS relative to maSS and selected miSS that are expressed at significantly high level at the given maSS expression level, separately in each tissue. Since the number of reads often exhibits an excess of zeros, we treated the total number of reads supporting a miSS (*r*_*min*_) in each tissue as a zero-inflated Poisson random variable with the parameters 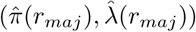 that depend on the number of reads supporting the corresponding maSS (*r*_*maj*_) as follows:

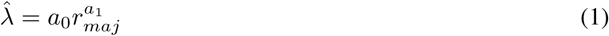

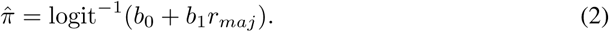

We estimated the parameters *a*_0_, *a*_1_, *b*_0_, and *b*_1_ separately in each tissue using zero-inflated Poisson (ZIP) regression model [68], computed the expected value of *r*_*min*_ for each miSS given the value of *r*_*maj*_, and assigned a P-value for each miSS as follows:

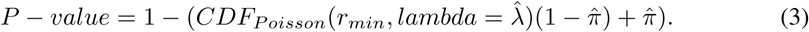

To account for multiple testing, we converted P-values to the false discovery rate, as measured by Q-value [38]. A miSS was called significantly expressed (or shortly significant) if it had the q-value below 5% in at least one tissue.

#### Tissue-specific miSS

The level of expression of a miSS relative to its corresponding maSS is reflected by the *φ* metric. To identify tissue-specific miSS among significantly expressed miSS, we analyzed the variability of the *φ* metric between and within tissues using the following linear regression model. For each significant miSS individually, we model *r*_*min*_ as a function of *r*_*maj*_ by the equation

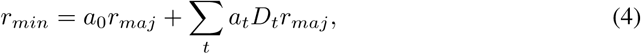

where *D*_*t*_ is a dummy variable corresponding to the tissue *t*. The slope *a*_*t*_ in this model can be interpreted as the change of the miSS relative usage in tissue *t* with respect to the tissue average as follows

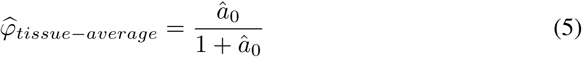

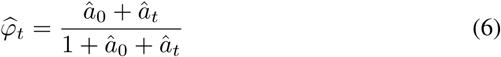

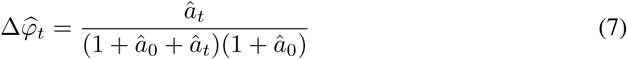

The significance of tissue-specific changes of *φ* represented by *a*_*t*_ can also be estimated using this linear model. This allows assigning P-values (and Q-values) to *a*_*t*_ for each miSS in each tissue. In order to filter out significant, but not substantial changes of tissue-specific miSS expression, we required the Q-value corresponding to *a*_*t*_ be below 5% and the absolute value of *a*_*t*_ be above 5%; a miSS satisfying these conditions was called tissue-specific in the tissue *t*. A miSS was called tissue-specific if it was specific in at least one tissue. Additionally, the sign of *a*_*t*_ allows to distinguish upregulation (*a*_*t*_ *>* 0) or downregulation (*a*_*t*_ *<* 0) of a miSS in the tissue *t*.

### Regulation of miSS by RBP

We used RNA-seq data from the experiments on the depletion of 248 RBPs in two human cell lines, K562 and HepG2 [69]. Short reads were mapped to the human genome by the data provider using STAR aligner v2.4.0k [65]. The read support of splice sites was called by IPSA pipeline as before (see processing of GTEx data). Out of 248 RBPs, we left only those for which 8 samples were present: two KD and two control samples for each of the two cell lines. Additionally, we required the presence of at least one publicly available eCLIP experiment [45] for each RBP. This confined our scope to 103 RBPs (S4 Table).

Out of 2,014 tissue-specific miSS, we selected 1,693 miSS that are supported by split reads in the KD or control samples of at least one RBP. The maSS were required to be stably expressed in both cell lines resulting in 1,308 tissue-specific miSS left.

For each miSS in each cell line, we constructed the following linear model:

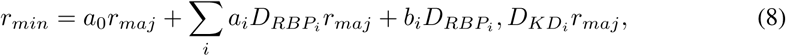

where 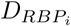 is a dummy variable corresponding to the RBP *i* and 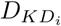 is a dummy variable corresponding to the knockdown of RBP *i* vs. control. The slope *b*_*i*_ in this model can be interpreted as the change of the miSS relative usage in the KD of RBP *i* compared to the control as follows:

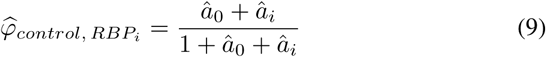

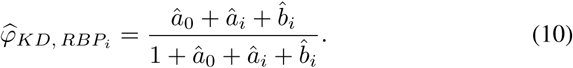

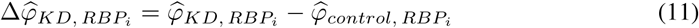

The significance of the *φ* change associated with the KD of RBP *i* allows assigning P-values and Q-values to *b*_*i*_ for each RBP-miSS pair in each cell line. We required the Q-value corresponding to *b*_*i*_ be below 5% in both HepG2 and K562 cell lines. In order to filter out significant, but not substantial changes of miSS expression, we additionally required that the absolute value of *b*_*i*_ be above 0.05. As a result, we obtained 168 significant RBP-miSS pairs, of which 24 pairs (14%) showed a discrepancy in the response to KD between cell lines (S9 Fig). These cases were excluded, and 144 RBP-miSS pairs (109 pairs with Δ*φ*_*KD*_ *>* 0 and 35 pairs with Δ*φ*_*KD*_ *<* 0) remained.

The gene read counts data was downloaded from GTEx (v7) portal on 08/05/2020 [70] and processed by DESeq2 package using apeglm shrinkage correction [71]. Differential expression analysis was done for each tissue against all other tissues. The p-values for 103 RBPs were Bonferroni-adjusted. An RBP was classified as tissue-specific if the adjusted p-value in the corresponding tissue was below 5% and the absolute value of log_2_ fold change was higher than 0.5. A tissue-specific RBP was considered upregulated in tissue *t* (Δ*RBP*_*t*_ *>* 0) if the log_2_ fold change value was positive and downregulated (Δ*RBP*_*t*_ *<* 0) otherwise.

The eCLIP peaks, which were called from the raw data by the producers, were downloaded from ENCODE data repository in bed format [69, 72]. The peaks in two immortalized human cell lines, K562 and HepG2, were filtered by the condition log FC ≥ 3 and P-value <0.001 as recommended [45]. Since the agreement between peaks in the two replicates was moderate (the median Jaccard distance 25% and 28% in K562 and HepG2, respectively), we took the union of peaks between the two replicates in two cell lines, and then pooled the resulting peaks.The presence of eCLIP peaks was assessed in the ±20 nt vicinity of a miSS position.

### Evidence of miSS translation in Ribo-Seq data

The global aggregate track of Ribo-Seq profiling, which tabulates the total number of footprint reads that align to the A-site of the elongating ribosome, was downloaded in bigWig format from GWIPS-viz Ribo-Seq genome browser [73]. It was intersected with TASS coordinates to obtain position-wise Ribo-Seq signal for miSS and maSS. The analysis was carried out on intronic miSS in TASS clusters of size 2. For each miSS, the relative Ribo-Seq support was calculated as

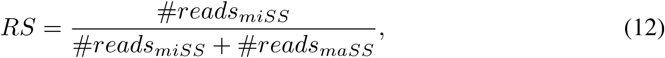

where #*reads*_*miSS*_ and #*reads*_*maSS*_ are the number of Ribo-Seq reads supporting the first exonic nucleotide of miSS and maSS, respectively. Higher values of *RS* indicate stronger evidence of translation.

### Structural annotation of miSS

All amino acids that are lost or gained due to using miSS instead of maSS were structurally annotated with respect to their spatial location in protein three-dimensional structure using StructMAn [74]. As a control, we also annotated all amino acids in all isoforms of the human proteome. Briefly, the procedure of structural annotation consists in mapping a particular amino acid into all experimentally resolved three-dimensional structures of proteins homologous to a given human isoform. The mapping is done by means of pairwise alignment of the respective protein sequences. Then the spatial location of the corresponding amino acid residue in the structure is analyzed in terms of proximity to other interaction partners (other proteins, nucleic acids, ligands, metal ions) and propensity to be exposed to the solvent or be buried in the protein core. Such annotations from different homologous proteins are then combined taking into account sequence similarity between the query human isoform and the proteins with the resolved structures, alignment coverage and the quality of the experimental structure. This resulted in structural annotations for 23,095,050 amino acids from 88,573 protein isoforms.

To use the structural annotation of amino acids in the analysis of TASS, we established a correspondence between 86,647 UniProt protein identifiers and 106,403 ENSEMBL transcripts identifiers discarding 3,194 transcripts that had ambiguous mappings [75].We used custom scripts to map 23,095,050 amino acids within structural annotation of UniProt entries to the human transcriptome and, furthermore, to the human genome using ENSEMBL transcript annotation. This procedure yielded 17,093,614 non-redundant genomic positions since some UniProt entries correspond to alternative isoforms of the same protein, and thus some amino acids from different entries can map to the same nucleotide in the genome. At that, positions that had ambiguous structural annotation from different transcripts were discarded.

Unlike maSS and exonic miSS, most of the intronic miSS are located outside of ENSEMBL transcripts and thus can not be directly classified based on the structural annotation. However, the structural annotation of exonic miSS coincides with that of the respective maSS in most cases (S10 Fig,A). We therefore assumed that the short distance between maSS and miSS allows to assign the structural annotation of the first exonic nucleotide of a maSS to all miSS including miSS located in introns. This way we defined structural annotation for 8,735 out of 16,327 frame-preserving expressed miSS in coding regions.

SLiM protein coordinates were mapped to genomic coordinates as described above. Regions between maSS and miSS (miSS indels) are compared with nearby exonic regions defined as the regions of the same length as miSS indels but located in the adjacent exons on the distance equal to the indel length. A SLiM is recognized to overlap with a particular region (miSS indel of nearby exonic region) if its genomic projection overlaps at least one exonic nucleotide.

### Evolutionary selection of miSS

Splice sites of annotated human transcripts were extracted from the comprehensive annotation of the human transcriptome (GENCODE v19 and NCBI RefSeq) using custom scripts [27, 28]. Internal boundaries of non-terminal exons (excluding splice sites overlapping with TASS clusters) were classified as constitutive splice sites if they were used as splice sites in all annotated transcripts. Position weight matrices were used to build consensus sequences for donor and acceptor constitutive splice sites as described in [76, 77]. Orthologs of the annotated human splice sites were identified in multiple sequence alignment of 46 vertebrate genomes with the human genome (GRCh37), which were downloaded from the UCSC Genome Browser in MAF format [64]. The alignments with marmoset and galago (bushbaby) genomes were extracted from MAF, and the alignment blocks were concatenated. The genomic sequence of splice sites in the common ancestor of human and marmoset with galago as an outgroup was reconstructed by parsimony [78]. Only splice sites with the canonical GT/AG dinucleotides in all three genomes were considered. The analysis was further confined to TASS clusters of size 2, in which only intronic miSS were considered to avoid the confounding effect of selection acting on the coding sequence in exonic miSS. This procedure resulted in 20,269 TASS (14,631 maSS and 5,638 miSS) in the coding regions and 3,586 TASS (2,111 maSS and 1,475 miSS) in the non-coding regions.

To estimate the strength of evolutionary selection acting on Cn and Nc nucleotides, we used a previously developed method with several modifications [61]. First, only intronic positions from the positions +3 to +6 for the donor splice sites and positions from -24 to -3 for the acceptor splice sites were considered (the canonical GT/AG dinucleotides were excluded as they were required to be conserved). The substitution counts were summed over all positions in these ranges. Furthermore, splice sites from the human genome were mapped onto the ancestral genome using MAF alignments but the substitutions were analyzed in the marmoset lineage, where the substitutions process goes independently from the human lineage (S11 Fig,B). This approach mitigates the systematic underrepresentation of Cn-to-Nc substitutions and the overrepresentation of Nc-to-Nc substitutions in the human lineage leading to artificial signs of strong positive and negative selection in cryptic and not significant miSS (S11 Fig,D) [61]. Constitutive splice sites were matched to maSS (miSS) by the ancestral strength using random sampling from the set of constitutive splice sites without replacement and requiring the strength difference not larger than 0.01.

### Mixture model for the estimation of the fraction of noisy miSS

The mixture model to estimate the fraction of noisy splice sites (Fig. 4C) was constructed as follows. Denote by *k* the size of the sample of interest (significantly expressed miSS, non-significantly expressed miSS, or maSS). We assume that the sample of interest is a mixture of two subsamples, *αk* splice sites from the negative set (cryptic splice sites, which demonstrate no evidence of selection, S11 Fig,D) and (1 − *α*)*k* splice sites from the positive set (all constitutive splice sites). For every *α* in the range from 0 to 1 with the step 0.0033, we sample randomly *αk* elements from the negative set and (1 − *α*)*k* elements from the positive set 300 times and construct the joint frequency distribution of *α* and *O/E*. To obtain the marginal (conditional) distribution corresponding to the observed value of *O/E* in the actual set of interest, we use an infinitesimal margin *E* to compute the empirical probability density in (*O/E* − *∈, O/E* + *∈*), and take the limit *∈* → 0 using the linear regression model *p* = *β*_0_ + *β*1*∈*. The quantiles were calculated for every *∈* in the range from 0.025 to 0.5 with the step 0.005 (S11 Fig,G). The interval estimates of *α* are inferred from the 2.5% and 97.5% quantiles.

### Statistical analysis

The data were analyzed and visualized using R statistics software version 3.4.1 and ggplot2 package, python version 2.7.5 and seaborn package. Non-parametric tests were performed with the statsmodels package using normal approximation with continuity correction. MW denotes Mann-Whitney sum of ranks test. Error bars in all figures and the numbers after the ±sign represent 95% confidence intervals. One-tailed P-values are reported throughout the paper, with the exception of linear regression models, in which we use two-tailed tests.

## Acknowledgments

AM and SD were supported by the Skolkovo Institute of Science and Technology Research Grant RF-0000000653 and Russian Foundation for Basic Research grant 18-29-13020-MK. AG acknowledges financial support from BMBF grant Sys_CARE (nr. 01ZX1908A) of the Federal German Ministry of Research and Education. All authors thank Drs. Sergei Moshkovskii and Mikhail Gorshkov and their research team for insightful discussions.

## Author contributions

AM and DP designed and carried out the statistical analysis. AG and OK designed and carried out structural annotation of the human proteome by homology modelling. SD designed and performed the evolutionary analysis. All authors discussed the results. AM, OK, and DP wrote the paper.

## Supporting information

**S1 Fig. The number of annotated**, *de novo* **and cryptic TASS in coding and non-coding regions**.

**S2 Fig. The intersection of TASS catalogue with TASSDB2**. Only a minor fraction of TASS in TASSDB2 are expressed.

**S3 Fig. The definition of the** *φ* **value** The definition of the *φ* value exemplified. A hypothetical maSS is supported by 4 split reads, while a hypothetical miSS is supported by 3 split reads, resulting in the *φ* value of 3*/*7.

**S4 Fig. TASS clusters of size three**. A TASS cluster of size three is characterized by two shift values: the rank two miSS relative to maSS, and rank three miSS relative to maSS. The top panel shows the joint distribution of rank two miSS shift (x-axis) and rank three miSS shift (y-axis) for donors (left) and (acceptors). The bottom panel shows LOGO charts of miSS sequences corresponding to shifts of +4 and -2 for the donor splice site, and +3 and +6 shifts for the acceptor splice site.

**S5 Fig. The distribution of shifts in coding vs. non-coding regions**. See Figure 1,E for comparison.

**S6 Fig. The strength of the consensus sequence of TASS. (A)** According to MaxEnt scores, maSS are on average stronger than miSS. **(B)** The relative usage of a miSS (*φ*) generally increases with increasing ΔMaxEnt value, its strength relative to that of the maSS. **(C)** The distribution of ΔMaxEnt values at different shifts in coding and non-coding regions. **(D)** The relative usage of a miSS (*φ*) as a function of the difference of TASS strengths. miSS with negative (upstream) shifts are used more frequently when the splice sites are nearly of the same strength.

**S7 Fig. Features of significantly expressed and tissue-specific miSS. (A)** The distribution of ΔMaxEnt values for miSS in different expression categories (left). The distribution of *RS* (RiboSeq support) values for miSS of different expression categories in protein-coding regions (right). **(B)** The average PhastCons scores (100 vertebrates) for positions near the miSS in different expression categories. **(C)** The distribution of PhastCons scores (100 vertebrates) at the consensus dinucleotides of splice sites (left) and at random dinucleotides in 30 nts intronic regions (right).

**S8 Fig. NAGNAGs and GYNNGYs. (A)** The intersection of the acceptor miSS located ±3 nts from the maSS with the list of NAGNAGs provided by Bradley et al [5]. **(B)** A NAGNAG acceptor splice site in the exon 20 of the MYRF gene. The upstream NAG is upregulated in the stomach, uterus, adipose tissues and downregulated in the brain. **(C)** The intersection of the donor miSS located ±4 nts from maSS with the list of GYNNGYs provided by Wang et al [1]. **(D)** A GYNNGY donor splice site in the exon 2 of the PAXX gene. The downstream GY is upregulated in the brain and downregulated in the stomach, pancreas, and liver tissues.

**S9 Fig. The response of a miSS to inactivation of an RBP in HepG2 (x-axis) and HepG2 (y-axis) cell lines**. Fractions of significant miSS-RBP pairs located in each quadrant are shown (the fractions are summed to 100

**S10 Fig. Structural annotation of miSS. (A)** The comparison of the structural annotation assigned directly to miSS (left) or from the structural annotation of the corresponding maSS (right). Only exonic miSS and corresponding maSS are considered. **(B)** The structural annotation for different categories of miSS. **(C)** Representation of short linear motifs (SLiMs) predicted with ELM server prediction tool in miSS of different expression categories. Only frame-preserving miSS in disordered coding regions are considered.

**S11 Fig. Evolutionary selection of miSS. (A)** The definition of the consensus (Cn) and non-consensus (Nc) nucleotide variants in the donor splice site. The definition for acceptor splice site is similar. **(B)** The evolutionary tree used to reconstruct the ancestral sequence of human and marmoset. **(C)** The computation of *obs* and *exp* statistics. **(D)** The selection of cryptic and not significant miSS in coding regions for marmoset and human genomes. **(E)** The selection of miSS in non-coding regions. **(F)** The distribution of ancestral strength for different splice site categories. **(G)** Estimation of the 95% confidence interval of *α* for different expression categories of miSS.

**S12 Fig. The dependence of the fraction of identified TASS on the number of considered samples**.

**S13 Fig. An example snapshot of the representation of the comprehensive catalogue of human TASS by Genome Browser track hub**.

**S1 Table. Summary statistics at different filtration steps of the TASS catalogue**.

**S2 Table. Characteristics of miSS in different expression categories. S3 Table. Abundance of tissue-specific miSS in tissues**.

**S4 Table. Accession codes for samples of shRNA RNP KD and eCLIP**.

**S5 Table. Predicted cases of miSS regulation by RBP with eCLIP support. S6 Table. Expressed miSS**.

## References

1. Wang ET, Sandberg R, Luo S, Khrebtukova I, Zhang L, Mayr C, et al. Alternative isoform regulation in human tissue transcriptomes. Nature. 2008;456(7221):470–476.

2. Raj B, Blencowe BJ. Alternative Splicing in the Mammalian Nervous System: Recent Insights into Mechanisms and Functional Roles. Neuron. 2015;87(1):14–27.

3. Merkin J, Russell C, Chen P, Burge CB. Evolutionary dynamics of gene and isoform regulation in Mammalian tissues. Science. 2012;338(6114):1593–1599.

4. Hiller M, Platzer M. Widespread and subtle: alternative splicing at short-distance tandem sites. Trends Genet. 2008;24(5):246–255.

5. Bradley RK, Merkin J, Lambert NJ, Burge CB. Alternative splicing of RNA triplets is often regulated and accelerates proteome evolution. PLoS Biol. 2012;10(1):e1001229.

6. Kozmik Z, Czerny T, Busslinger M. Alternatively spliced insertions in the paired domain restrict the DNA sequence specificity of Pax6 and Pax8. EMBO J. 1997;16(22):6793–6803.

7. Tadokoro K, Yamazaki-Inoue M, Tachibana M, Fujishiro M, Nagao K, Toyoda M, et al. Frequent occurrence of protein isoforms with or without a single amino acid residue by subtle alternative splicing: the case of Gln in DRPLA affects subcellular localization of the products. J Hum Genet. 2005;50(8):382–394.

8. Yan M, Wang LC, Hymowitz SG, Schilbach S, Lee J, Goddard A, et al. Two-amino acid molecular switch in an epithelial morphogen that regulates binding to two distinct receptors. Science. 2000;290(5491):523–527.

9. Mullaney JM, Mills RE, Pittard WS, Devine SE. Small insertions and deletions (INDELs) in human genomes. Hum Mol Genet. 2010;19(R2):R131–136.

10. Auton A, Brooks LD, Durbin RM, Garrison EP, Kang HM, Korbel JO, et al. A global reference for human genetic variation. Nature. 2015;526(7571):68–74.

11. Collins FS, Drumm ML, Cole JL, Lockwood WK, Vande Woude GF, Iannuzzi MC. Construction of a general human chromosome jumping library, with application to cystic fibrosis. Science. 1987;235(4792):1046–1049.

12. Irimia M, Weatheritt RJ, Ellis JD, Parikshak NN, Gonatopoulos-Pournatzis T, Babor M, et al. A highly conserved program of neuronal microexons is misregulated in autistic brains. Cell. 2014;159(7):1511–1523.

13. Lin M, Whitmire S, Chen J, Farrel A, Shi X, Guo JT. Effects of short indels on protein structure and function in human genomes. Sci Rep. 2017;7(1):9313.

14. Hiller M, Szafranski K, Huse K, Backofen R, Platzer M. Selection against tandem splice sites affecting structured protein regions. BMC Evol Biol. 2008;8:89.

15. Hiller M, Huse K, Szafranski K, Jahn N, Hampe J, Schreiber S, et al. Widespread occurrence of alternative splicing at NAGNAG acceptors contributes to proteome plasticity. Nat Genet. 2004;36(12):1255–1257.

16. Sinha R, Nikolajewa S, Szafranski K, Hiller M, Jahn N, Huse K, et al. Accurate prediction of NAGNAG alternative splicing. Nucleic Acids Res. 2009;37(11):3569–3579.

17. Wang M, Zhang P, Shu Y, Yuan F, Zhang Y, Zhou Y, et al. Alternative splicing at GYNNGY 5’ splice sites: more noise, less regulation. Nucleic Acids Res. 2014;42(22):13969–13980.

18. Tsai KW, Chan WC, Hsu CN, Lin WC. Sequence features involved in the mechanism of 3’ splice junction wobbling. BMC Mol Biol. 2010;11:34.

19. Chern TM, van Nimwegen E, Kai C, Kawai J, Carninci P, Hayashizaki Y, et al. A simple physical model predicts small exon length variations. PLoS Genet. 2006;2(4):e45.

20. Dou Y, Fox-Walsh KL, Baldi PF, Hertel KJ. Genomic splice-site analysis reveals frequent alternative splicing close to the dominant splice site. RNA. 2006;12(12):2047–2056.

21. Tress ML, Abascal F, Valencia A. Alternative Splicing May Not Be the Key to Proteome Complexity. Trends Biochem Sci. 2017;42(2):98–110.

22. Tress ML, Abascal F, Valencia A. Most Alternative Isoforms Are Not Functionally Important. Trends Biochem Sci. 2017;42(6):408–410.

23. Blencowe BJ. The Relationship between Alternative Splicing and Proteomic Complexity. Trends Biochem Sci. 2017;42(6):407–408.

24. Sinha R, Lenser T, Jahn N, Gausmann U, Friedel S, Szafranski K, et al. TassDB2 - A comprehensive database of subtle alternative splicing events. BMC Bioinformatics. 2010;11:216.

25. Pan Q, Shai O, Lee LJ, Frey BJ, Blencowe BJ. Deep surveying of alternative splicing complexity in the human transcriptome by high-throughput sequencing. Nat Genet. 2008;40(12):1413–1415.

26. Melé M, Ferreira PG, Reverter F, DeLuca DS, Monlong J, Sammeth M, et al. Human genomics. The human transcriptome across tissues and individuals. Science. 2015;348(6235):660–665.

27. Harrow J, Frankish A, Gonzalez JM, Tapanari E, Diekhans M, Kokocinski F, et al. GENCODE: the reference human genome annotation for The ENCODE Project. Genome Res. 2012;22(9):1760–1774.

28. O’Leary NA, Wright MW, Brister JR, Ciufo S, Haddad D, McVeigh R, et al. Reference sequence (RefSeq) database at NCBI: current status, taxonomic expansion, and functional annotation. Nucleic Acids Res. 2016;44(D1):D733–745.

29. Bretschneider H, Gandhi S, Deshwar AG, Zuberi K, Frey BJ. COSSMO: predicting competitive alternative splice site selection using deep learning. Bioinformatics. 2018;34(13):i429–i437.

30. Nellore A, Jaffe AE, Fortin JP, Alquicira-Hernández J, Collado-Torres L, Wang S, et al. Human splicing diversity and the extent of unannotated splice junctions across human RNA-seq samples on the Sequence Read Archive. Genome Biol. 2016;17(1):266.

31. Pervouchine DD, Knowles DG, Guigó R. Intron-centric estimation of alternative splicing from RNA-seq data. Bioinformatics. 2013;29(2):273–274.

32. Pickrell JK, Pai AA, Gilad Y, Pritchard JK. Noisy splicing drives mRNA isoform diversity in human cells. PLoS Genet. 2010;6(12):e1001236.

33. Busch A, Hertel KJ. Extensive regulation of NAGNAG alternative splicing: new tricks for the spliceosome? Genome Biol. 2012;13(2):143.

34. Lykke-Andersen S, Chen Y, Ardal BR, Lilje B, Waage J, Sandelin A, et al. Human nonsense-mediated RNA decay initiates widely by endonucleolysis and targets snoRNA host genes. Genes Dev. 2014;28(22):2498–2517.

35. Chua K, Reed R. An upstream AG determines whether a downstream AG is selected during catalytic step II of splicing. Mol Cell Biol. 2001;21(5):1509–1514.

36. Barash Y, Calarco JA, Gao W, Pan Q, Wang X, Shai O, et al. Deciphering the splicing code. Nature. 2010;465(7294):53–59.

37. Saudemont B, Popa A, Parmley JL, Rocher V, Blugeon C, Necsulea A, et al. The fitness cost of mis-splicing is the main determinant of alternative splicing patterns. Genome Biol. 2017;18(1):208.

38. Storey JD, Tibshirani R. Statistical significance for genomewide studies. Proc Natl Acad Sci USA. 2003;100(16):9440–9445.

39. Gong D, Chi X, Ren K, Huang G, Zhou G, Yan N, et al. Structure of the human plasma membrane Ca2+-ATPase 1 in complex with its obligatory subunit neuroplastin. Nat Commun. 2018;9(1):3623.

40. Beesley PW, Herrera-Molina R, Smalla KH, Seidenbecher C. The Neuroplastin adhesion molecules: key regulators of neuronal plasticity and synaptic function. J Neurochem. 2014;131(3):268–283.

41. Xu Q, Modrek B, Lee C. Genome-wide detection of tissue-specific alternative splicing in the human transcriptome. Nucleic Acids Res. 2002;30(17):3754–3766.

42. Craxton A, Munnur D, Jukes-Jones R, Skalka G, Langlais C, Cain K, et al. PAXX and its paralogs synergistically direct DNA polymerase Î” activity in DNA repair. Nat Commun. 2018;9(1):3877.

43. Grosso AR, Gomes AQ, Barbosa-Morais NL, Caldeira S, Thorne NP, Grech G, et al. Tissue-specific splicing factor gene expression signatures. Nucleic Acids Res. 2008;36(15):4823–4832.

44. Nostrand ELV, Freese P, Pratt GA, Wang X, Wei X, Blue SM, et al. A Large-Scale Binding and Functional Map of Human RNA Binding Proteins. bioRxiv. 2017;doi: 10.1101/179648.

45. Van Nostrand EL, Pratt GA, Shishkin AA, Gelboin-Burkhart C, Fang MY, Sundararaman B, et al. Robust transcriptome-wide discovery of RNA-binding protein binding sites with enhanced CLIP (eCLIP). Nat Methods. 2016;13(6):508–514.

46. Hall MP, Nagel RJ, Fagg WS, Shiue L, Cline MS, Perriman RJ, et al. Quaking and PTB control overlapping splicing regulatory networks during muscle cell differentiation. RNA. 2013;19(5):627–638.

47. Romero PR, Zaidi S, Fang YY, Uversky VN, Radivojac P, Oldfield CJ, et al. Alternative splicing in concert with protein intrinsic disorder enables increased functional diversity in multicellular organisms. Proc Natl Acad Sci USA. 2006;103(22):8390–8395.

48. Davey NE, Van Roey K, Weatheritt RJ, Toedt G, Uyar B, Altenberg B, et al. Attributes of short linear motifs. Mol Biosyst. 2012;8(1):268–281.

49. Van Roey K, Uyar B, Weatheritt RJ, Dinkel H, Seiler M, Budd A, et al. Short linear motifs: ubiquitous and functionally diverse protein interaction modules directing cell regulation. Chem Rev. 2014;114(13):6733–6778.

50. Uyar B, Weatheritt RJ, Dinkel H, Davey NE, Gibson TJ. Proteome-wide analysis of human disease mutations in short linear motifs: neglected players in cancer? Mol Biosyst. 2014;10(10):2626–2642.

51. Huang KY, Lee TY, Kao HJ, Ma CT, Lee CC, Lin TH, et al. dbPTM in 2019: exploring disease association and cross-talk of post-translational modifications. Nucleic Acids Res. 2019;47(D1):D298–D308.

52. Tian Y, Chang JC, Fan EY, Flajolet M, Greengard P. Adaptor complex AP2/PICALM, through interaction with LC3, targets Alzheimer’s APP-CTF for terminal degradation via autophagy. Proc Natl Acad Sci USA. 2013;110(42):17071–17076.

53. Moreau K, Fleming A, Imarisio S, Lopez Ramirez A, Mercer JL, Jimenez-Sanchez M, et al. PICALM modulates autophagy activity and tau accumulation. Nat Commun. 2014;5:4998.

54. Johansen T, Lamark T. Selective Autophagy: ATG8 Family Proteins, LIR Motifs and Cargo Receptors. J Mol Biol. 2020;432(1):80–103.

55. Wang X, Zamore PD, Hall TM. Crystal structure of a Pumilio homology domain. Mol Cell. 2001;7(4):855–865.

56. Yang J, Zhang Y. I-TASSER server: new development for protein structure and function predictions. Nucleic Acids Res. 2015;43(W1):W174–181.

57. Wood CW, Ibarra AA, Bartlett GJ, Wilson AJ, Woolfson DN, Sessions RB. BAlaS: fast, interactive and accessible computational alanine-scanning using BudeAlaScan. Bioinformatics. 2020;36(9):2917–2919.

58. Bobo-Jiménez V, Delgado-Esteban M, Angibaud J, Sánchez-Morán I, de la Fuente A, Yajeya J, et al. APC/CCdh1-Rock2 pathway controls dendritic integrity and memory. Proc Natl Acad Sci USA. 2017;114(17):4513–4518.

59. Delgado-Esteban M, García-Higuera I, Maestre C, Moreno S, Almeida A. APC/C-Cdh1 coordinates neurogenesis and cortical size during development. Nat Commun. 2013;4:2879.

60. Raney BJ, Dreszer TR, Barber GP, Clawson H, Fujita PA, Wang T, et al. Track data hubs enable visualization of user-defined genome-wide annotations on the UCSC Genome Browser. Bioinformatics. 2014;30(7):1003–1005.

61. Denisov SV, Bazykin GA, Sutormin R, Favorov AV, Mironov AA, Gelfand MS, et al. Weak negative and positive selection and the drift load at splice sites. Genome Biol Evol. 2014;6(6):1437–1447.

62. Irimia M, Roy SW, Neafsey DE, Abril JF, Garcia-Fernandez J, Koonin EV. Complex selection on 5’ splice sites in intron-rich organisms. Genome Res. 2009;19(11):2021–2027.

63. Razeto-Barry P, Díaz J, Vásquez RA. The nearly neutral and selection theories of molecular evolution under the fisher geometrical framework: substitution rate, population size, and complexity. Genetics. 2012;191(2):523–534.

64. Haeussler M, Zweig AS, Tyner C, Speir ML, Rosenbloom KR, Raney BJ, et al. The UCSC Genome Browser database: 2019 update. Nucleic Acids Res. 2019;47(D1):D853–D858.

65. Dobin A, Davis CA, Schlesinger F, Drenkow J, Zaleski C, Jha S, et al. STAR: ultrafast universal RNA-seq aligner. Bioinformatics. 2013;29(1):15–21.

66. Yeo G, Burge CB. Maximum entropy modeling of short sequence motifs with applications to RNA splicing signals. J Comput Biol. 2004;11(2-3):377–394.

67. Wang L, Wang S, Li W. RSeQC: quality control of RNA-seq experiments. Bioinformatics. 2012;28(16):2184–2185.

68. Zeileis A, Kleiber C, Jackman S. Regression Models for Count Data inR. Journal of Statistical Software. 2008;27(8). doi: 10.18637/jss.v027.i08.

69. Dunham I, Kundaje A, Aldred SF, Collins PJ, Davis CA, Doyle F, et al. An integrated encyclopedia of DNA elements in the human genome. Nature. 2012;489(7414):57–74.

70. Lonsdale J, Thomas J, Salvatore M, Phillips R, Lo E, Shad S, et al. The Genotype-Tissue Expression (GTEx) project. Nat Genet. 2013;45(6):580–585.

71. Zhu A, Ibrahim JG, Love MI. Heavy-tailed prior distributions for sequence count data: removing the noise and preserving large differences. Bioinformatics. 2019;35(12):2084–2092.

72. Sloan CA, Chan ET, Davidson JM, Malladi VS, Strattan JS, Hitz BC, et al. ENCODE data at the ENCODE portal. Nucleic Acids Res. 2016;44(D1):D726–732.

73. Michel AM, Fox G, Mkiran A, De Bo C, O’Connor PB, Heaphy SM, et al. GWIPS-viz: development of a ribo-seq genome browser. Nucleic Acids Res. 2014;42(Database issue):D859–864.

74. Gress A, Ramensky V, Büch J, Keller A, Kalinina OV. StructMAn: annotation of single-nucleotide polymorphisms in the structural context. Nucleic Acids Res. 2016;44(W1):W463–468.

75. authors listed N. UniProt: a worldwide hub of protein knowledge. Nucleic Acids Res. 2019;47(D1):D506–D515.

76. Stamm S, Zhu J, Nakai K, Stoilov P, Stoss O, Zhang MQ. An alternative-exon database and its statistical analysis. DNA Cell Biol. 2000;19(12):739–756.

77. Denisov S, Bazykin G, Favorov A, Mironov A, Gelfand M. Correlated Evolution of Nucleotide Positions within Splice Sites in Mammals. PLoS ONE. 2015;10(12):e0144388.

78. Farris JS. Methods for Computing Wagner Trees. Systematic Biology. 1970;19(1):83–92. doi: 10.1093/sysbio/19.1.83.

79. Park E, Pan Z, Zhang Z, Lin L, Xing Y. The Expanding Landscape of Alternative Splicing Variation in Human Populations. Am J Hum Genet. 2018;102(1):11–26.

80. Pervouchine D, Popov Y, Berry A, Borsari B, Frankish A, Guigó R. Integrative transcriptomic analysis suggests new autoregulatory splicing events coupled with nonsense-mediated mRNA decay. Nucleic Acids Res. 2019;47(10):5293–5306.

81. Ni JZ, Grate L, Donohue JP, Preston C, Nobida N, O’Brien G, et al. Ultraconserved elements are associated with homeostatic control of splicing regulators by alternative splicing and nonsense-mediated decay. Genes Dev. 2007;21(6):708–718.

82. Lareau LF, Brenner SE. Regulation of splicing factors by alternative splicing and NMD is conserved between kingdoms yet evolutionarily flexible. Mol Biol Evol. 2015;32(4):1072–1079.

83. Röst HL, Malmström L, Aebersold R. Reproducible quantitative proteotype data matrices for systems biology. Mol Biol Cell. 2015;26(22):3926–3931.

84. Wang X, Codreanu SG, Wen B, Li K, Chambers MC, Liebler DC, et al. Detection of Proteome Diversity Resulted from Alternative Splicing is Limited by Trypsin Cleavage Specificity. Mol Cell Proteomics. 2018;17(3):422–430.

85. Weatheritt RJ, Sterne-Weiler T, Blencowe BJ. The ribosome-engaged landscape of alternative splicing. Nat Struct Mol Biol. 2016;23(12):1117–1123.

86. Ellis JD, Barrios-Rodiles M, Colak R, Irimia M, Kim T, Calarco JA, et al. Tissue-specific alternative splicing remodels protein-protein interaction networks. Mol Cell. 2012;46(6):884–892.

87. Buljan M, Chalancon G, Dunker AK, Bateman A, Balaji S, Fuxreiter M, et al. Alternative splicing of intrinsically disordered regions and rewiring of protein interactions. Curr Opin Struct Biol. 2013;23(3):443–450.

